# Cryo-EM reconstruction of the human 40S ribosomal subunit at 2.15 Å resolution

**DOI:** 10.1101/2022.01.16.475527

**Authors:** Simone Pellegrino, Kyle C. Dent, Tobias Spikes, Alan J. Warren

**Author notes:** Present address: Medical Research Council Laboratory of Molecular Biology, Cambridge CB2 0QH, United Kingdom (Kyle C. Dent). To whom correspondence should be addressed (S.P.), (A.J.W.).

## Abstract

The chemical modification of ribosomal RNA and proteins is critical for ribosome assembly, for protein synthesis and may drive ribosome specialization in development and disease. However, the inability to accurately visualize these modifications has limited mechanistic understanding of the role of these modifications in ribosome function. Here we report the 2.15 Å resolution cryo-EM reconstruction of the human 40S ribosomal subunit. We directly visualize post-transcriptional modifications within the 18S rRNA and post-translational modifications at the N-termini of two ribosomal proteins. Additionally, we interpret the solvation shells in the core regions of the 40S ribosomal subunit and reveal how potassium and magnesium ions establish both universally conserved and eukaryote-specific coordination to promote the stabilization and folding of key ribosomal elements. This work provides unprecedented structural details for the human 40S ribosomal subunit that will serve as an important reference for unraveling the functional role of ribosomal RNA modifications.

## INTRODUCTION

The ribosome is a ribonucleoprotein machine that translates the genetic code in every living organism. It consists of two subunits, comprising 4 ribosomal RNAs (rRNAs) and 80 ribosomal proteins (RPs). The large (60S) subunit is responsible for peptide bond formation, while the small (40S) subunit reads and decodes the messenger RNA (mRNA). Assembly of ribosomal subunits is one of the most energy consuming processes within cells, requiring more than 300 biogenesis factors and 80 small nucleolar RNAs (snoRNAs) in humans (Tafforeau et al. 2013; Bohnsack and Bohnsack 2019). Similarly, protein synthesis is a tightly regulated process that involves the sequential action of translation factors to ensure speed and accuracy of the process (Melnikov et al. 2012).

The rRNAs and RPs are decorated with several types of modifications. Post-transcriptional modifications are mainly incorporated early in the process of ribosome biogenesis, likely soon after the primary 47S transcript is generated (Aubert et al. 2018; Bohnsack and Bohnsack 2019). There are 14 distinct rRNA modifications within the human 80S ribosome, which decorate approximately 5% of rRNA residues. These modifications are thought to stabilize rRNA structure and participate in the binding of functional ligands during ribosome biogenesis and translation (Natchiar et al. 2017; Bohnsack and Bohnsack 2019; Taoka et al. 2018). Eukaryotic rRNA modifications mainly comprise 2’-O-methylation of rRNA ribose and the isomerization of the nucleoside uridine to pseudouridine (psi, ψ); these chemical modifications are introduced by box C/D and box H/ACA snoRNPs, respectively, with nucleotide specificity dictated by the sequence complementarity of snoRNAs, which pair with the target site (Ojha et al. 2020; Sloan et al. 2017). All the other rRNA modifications involve only nucleotide bases and are introduced by stand-alone enzymes that function throughout the ribosome biogenesis pathway (Sloan et al. 2017). Distinct patterns of rRNA modifications may promote ribosome specialization, promoting the selective translation of subsets of mRNAs (Erales et al. 2017; Jansson et al. 2021; Pauli et al. 2020). Furthermore, rRNA modifications may also be dysregulated in cancer (Gong et al. 2017; Valleron et al. 2012a, 2012b; Heiss et al. 1998; Zhou et al. 2017; Barbieri and Kouzarides 2020; Monaco et al. 2018; Janin et al. 2020). Methods such as 2D thin layer chromatography, primer extension-based assays, or chemical derivatization were initially used to identify 2′-O-methylation and pseudouridylation sites (Maden 1986, 1988; Maden and Wakeman 1988; Maden et al. 1995). More recently, “stable isotope–labeled ribonucleic acid as an internal standard” (SILNAS)-based quantification of post-transcriptional modifications (Taoka et al. 2018) and cryo-electron microscopy (cryo-EM) (Natchiar et al. 2017) approaches have sought to identify the complete repertoire of modifications on the human 80S ribosome. However, given the discrepancies in the published data between these methods, higher resolution structures of human ribosomes are required to directly visualize rRNA modifications with greater accuracy to better resolve their function.

Here, we report a cryo-EM reconstruction of the human 40S ribosomal subunit at an overall resolution of 2.15 Å, extending to 1.8 Å in the core, revealing unprecedented structural details. We visualize post-transcriptional modifications incorporated into the 18S rRNA of the 40S ribosomal subunit and identify experimental density for post-translational modifications of two ribosomal proteins. Furthermore, we model the solvation shells of the 40S ribosomal subunit, revealing the position of monovalent and divalent cations.

## RESULTS

The small ribosomal subunit consists of two domains, the body and the head (**Fig. 1 A, Fig. S1**). The head is highly dynamic and rotates (“swivels”) with respect to the main axis of the body when not interacting with functional ligands, such as transfer RNA (tRNA), mRNA, assembly or translation factors. The intrinsic mobility of these two domains and the presence of elements that are flexible when not interacting with the large ribosomal subunit, such as h44, have hampered efforts to achieve high-resolution reconstructions of the 40S ribosomal subunit. By employing a novel method for sample preparation (see Materials and Methods) and by implementing the multibody refinement algorithm in Relion 3.0 (Nakane and Scheres 2021)(Nakane and Scheres 2021) (**Fig. S1 and S2**), we obtained individual reconstructions of both the head and body at nominal resolutions of 2.24 and 2.09 Å, respectively, extending to 1.8 Å resolution in the most stable regions (**Fig. 1 A**, **Table S1, Fig. S2**). Our cryo-EM reconstructions allowed us to visualize unambiguously 73 out of the 91 rRNA modifications of the human 40S ribosomal subunit identified by SILNAS-based quantification (Taoka et al. 2018) (**Table S2, Fig. S3 and S4**). We visualized chemical modifications that were either previously unassigned (Natchiar et al. 2017) or were yet unmodeled in the human 40S ribosomal subunit, including 2’-O-methylation (rRNA residues G1447 and A590) (**Fig. 1 B-C**), and uridine isomerization (residues U572 and U863) (**Fig. 1 D-E, Fig. S3 and S4**). Our findings are in good overall agreement with data from 2OMet-seq (Incarnato et al. 2017), RiboMeth-seq (Krogh et al. 2016), Pseudo-seq (Carlile et al. 2014) and SILNAS-based analysis of rRNA modifications (Taoka et al. 2018). Of the four new modifications identified by Taoka et al. (2’-O-methylation of residue C621 and pseudouridines ψ897, ψ1045, ψ1136 and ψ1232), we visualized three of the four proposed pseudouridines (ψ1045, ψ1136 and ψ1232), while ψ897 localizes to a flexible region where the cryo-EM density could not be interpreted (**Fig. 1 F, S3 and S4, Table S2**). While the imino group at position 1 of ψ1045 directly contacts the 2’-OH group of the neighboring G1044, the isomeric nucleobases of ψ1136 and ψ1242 establish water-mediated hydrogen bonds with the phosphate backbone. Interestingly, ψ1136 was present in low sub-stoichiometric amounts in human TK6 cells using SILNAS (Taoka et al. 2018), but was not identified in previous cryo-EM reconstructions of either the human 80S ribosome (Natchiar et al. 2017) or late pre-40S subunits (Ameismeier et al. 2020). However, our map revealed clear density for a water molecule mediating an interaction between the N1 imino group of uridine and the phosphate backbone (**Fig. 1 F and S4**). By contrast, we did not identify 2’-O-methylation of residue C621, a modification that would likely disrupt water-mediated hydrogen bonding interactions between the two nucleobases of residues C621 and G6 (**Fig. S4 A**). Both the map and the atomic model of a human late pre-40S particle (PDB ID: 6ZXG) further support the absence of 2’-O-methylation of residue C621.

**Figure 1:**
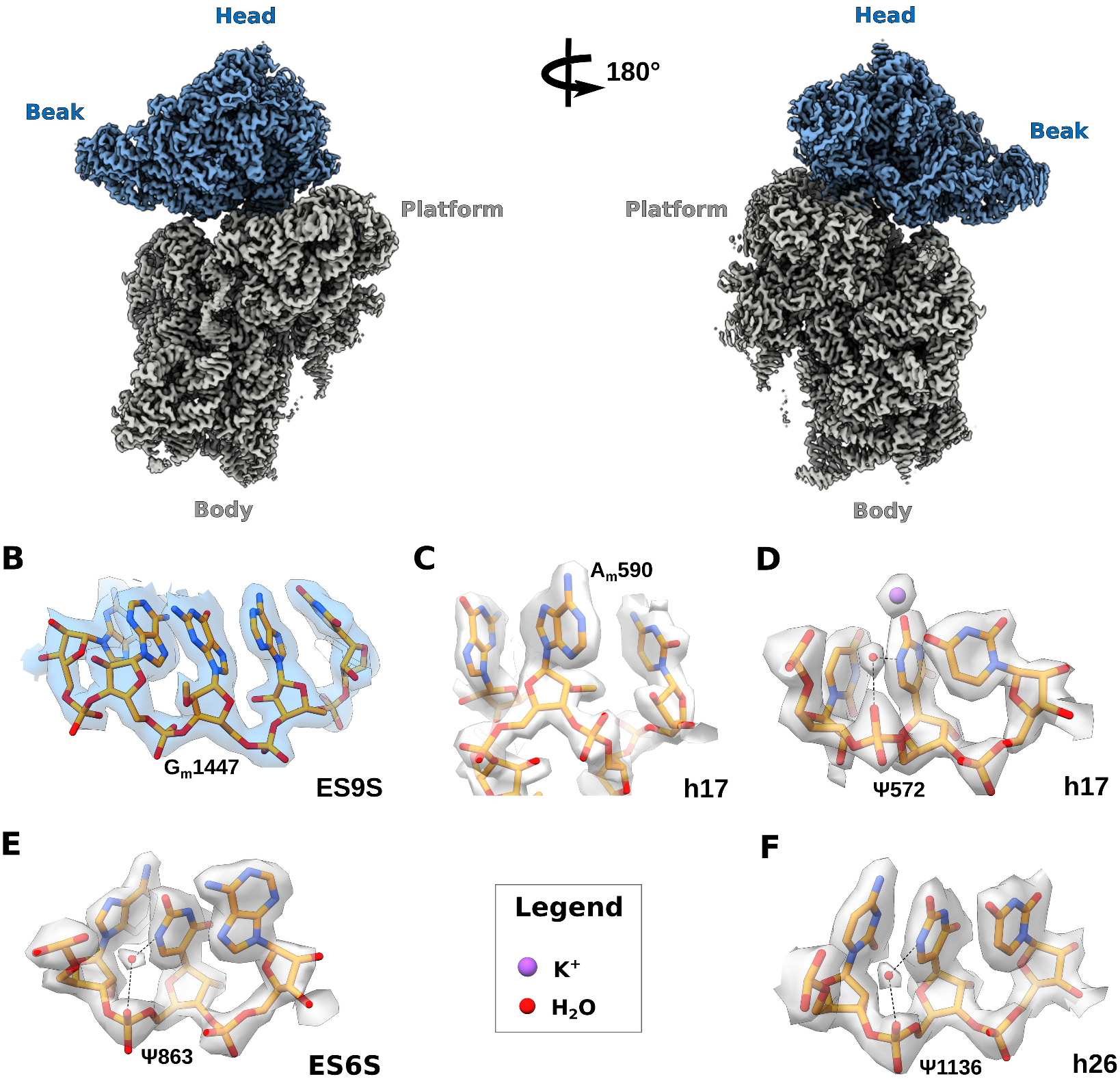
Newly visualized rRNA modifications of the human 40S subunit at 2.15 Å resolution. A) Cryo-EM reconstruction of the human 40S ribosomal subunit viewed from the intersubunit interface (left) and rotated 180° (right). The map corresponding to the head domain is colored blue, body in grey. Water molecules and K^+^ ions are shown as spheres (legend at the bottom of the panel). B) 2’-O-methylated G1447 in expansion segment 9S (ES9S) of h39. C) Close-up of the tip of h17, showing 2’-O-methylated A590. D) Uridine isomerization at residue U572 in h17. E) Uridine isomerization at residue U863, within ES6S. F) Uridine isomerization at residue U1136, with a water molecule coordinating the imino group at position 1.

Water molecules, monovalent and divalent cations are important for ribosome structure and function (Rozov et al. 2019; Klein et al. 2004; Watson et al. 2020). In our maps, we identified monovalent cations bound to both rRNA and RPs. We interpreted these as K^+^ ions given the composition of the buffers used during sample purification, together with their coordination geometry and distance. Interestingly, compared to the bacterial 30S ribosomal subunit (PDB ID: 6QNR) (Rozov et al. 2019), the binding position of monovalent cations is conserved in key regions of the human 40S subunit. For example, we identified a K^+^ ion bound between h42 and uS13, which participates in formation of the neck-head junction within the head domain by coordinating the phosphate backbone of U1631 with the carbonyl groups of the backbone of residues Thr31, Ile33 and Val36 (**Fig. 2 A**). Similarly, in bacteria the protein backbone of uS13 is engaged in K^+^-mediated coordination with rRNA residue U1330; however, as the carbonyl group of the Ile25 backbone in bacteria does not face the ion binding pocket, it is therefore unable to interact with the K^+^ (**Fig. S5 A**). Furthermore, in proximity to the modified residue m^7^G1639, we identified a K^+^ ion bound to the nucleobases of G1638, G1226 and G1227 and the phosphate backbone of G1226 with square anti-prismatic geometry (**Fig. 2 B**). An identical position for a K^+^ ion is found in bacteria (**Fig. S5 B**). Human residue G626 (G530 bacterial numbering) on h18 is involved in the decoding mechanism, switching from a *syn* conformation when the A-site is vacant, to the *anti* conformation when tRNA is bound (Ogle et al. 2001; Shao et al. 2016). Compared to the bacterial 70S ribosome (PDB: 6QNR), our model highlights a conserved K^+^ ion that interacts with residues C614 and G625 of h18, the hydroxyl group of the side chain of Asn63 and the carbonyl group of Pro62 of uS12 (**Fig. 2 C**). Supporting our findings, in bacteria a K^+^ ion has also been experimentally assigned in this conserved pocket (**Fig. S5 C)**. In our reconstructions, potassium ions mostly stabilize rRNA by coordinating stacked nucleotides, rather than by interacting with the phosphate backbone, consistent with data in bacteria (Rozov et al. 2019). For instance, within h28, a K^+^ ion is involved in extensive coordination with rRNA nucleobases, that likely maintains base planarity (**Fig. 2 D**). Residue ψ1692 forms a non-complementary Watson-Crick base pair with G1206 in which the ψ1692 oxygen atom at position 4 directly interacts with the K^+^ ion. In bacteria, while the conserved uridine is unmodified (U1391), a K^+^ ion nevertheless establishes a similar bonding pattern (**Fig. S5 D**).

**Figure 2:**
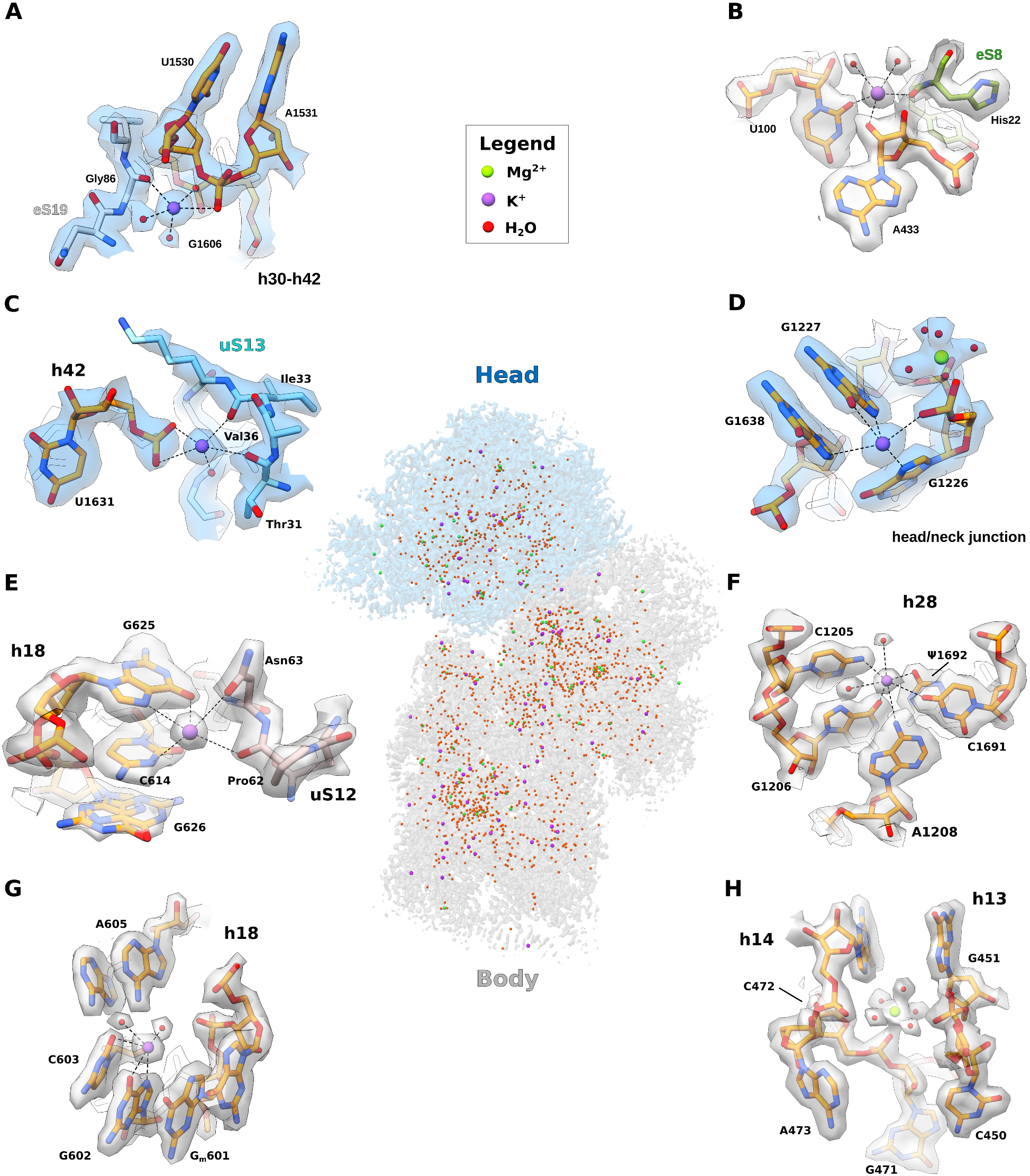
Solvation shell of the human 40S subunit. Middle of panel: map of the head (blue) and body (grey) domains; water molecules, K^+^ and Mg^2+^ ions are shown as spheres (legend at the top of the panel). A) K^+^ ion bridges the universally conserved ribosomal protein uS13 and h42 within the head domain of the 18S rRNA. B) K^+^ ion maintains rRNA planarity within the head/neck region close to h29. C) K^+^ ion stabilizes the h18-uS12 interaction within the decoding center. D) Detail of K^+^ ion and water coordination within h28. E) K^+^ ion bridges the eukaryote-specific ribosomal protein eS19 and the 18S rRNA. F) K^+^ ion bridging eukaryote-specific eS8 and 18S rRNA. G) Close-up of h18 within the body domain of the 18S rRNA showing detail of eukaryote-specific coordination of a K^+^ ion and water molecules with rRNA residues. H) Detail of the solvation shell of h13 and the proximal h14, including a Mg^2+^ ion with octahedral coordination.

In addition, we visualized K^+^ ions bound in eukaryote-specific positions. For example, we identified a K^+^ ion bound between the peptide backbone of the eukaryote-specific RP eS19 (residue Gly86) and the phosphate backbone of rRNA helices h30 and h42 (residues A1531 and G1606, respectively), which also interacts with two water molecules in square anti-prismatic coordination (**Fig. 2 E**). Similarly, in the three-way junction between helices h7, h11 and h12, a K^+^ ion bridged the 2’-OH group of A433, the oxygen at position 2 of U100 and the peptide backbone of eS8 (residue His22) (**Fig. 2F**). We identified a K^+^ ion bound to the nucleobases of G602 and C603 at the bulge of h18, consistent with a key role in promoting rRNA stacking. Additionally, we identified two water molecules that coordinate the interaction between the nucleobase of A605 and the phosphate backbone of G_m_601 (**Fig. 2 G**). The evolution of the ribosome may have driven the requirement for ions. For example, on h13, rRNA residue C450 (C330 in *T. thermophilus*) adopts a different conformation in eukaryotes (**Fig. 2 H, Fig. S5 E**), likely to allow access for the N-terminus of the eukaryotic specific ribosomal protein eS4. Interestingly, in the human map, a Mg^2+^ ion is bound with octahedral coordination to the phosphate backbone of C472, which establishes water-mediated contacts to the backbone and sugar moiety of G451 (**Fig. 2 H**). By contrast, in bacteria a K^+^ ion is found in this conserved position (**Fig. S5 E**).

Our cryo-EM reconstruction revealed the presence of additional density at the N-terminus of two ribosomal proteins, uS2 and eS21 (**Fig. 3 A and B**). uS2 and eS21 are part of the so-called S0-cluster and are among the last ribosomal proteins assembled into the 40S ribosomal subunit (Linnemann et al. 2019). Recruitment of the S0-cluster is thought to occur in the cytoplasm, after remodeling of the head domain and release of the biogenesis factor RRP12 (Linnemann et al. 2019; Ameismeier et al. 2018). We hypothesized that the additional density might represent post-translational modifications, consistent with mass spectrometry data and a recent cryo-EM reconstruction of 80S ribosomes purified from rabbit reticulocytes (van de Waterbeemd et al. 2018; Yu et al. 2005; Bhatt et al. 2021). We therefore interrogated our human 40S sample for post-translational modifications using liquid chromatography mass spectrometry (LC-MS/MS) (**Table S3**). Analysis of the tryptic peptides confirmed the presence of N-terminal acetylated uS2 on residue Ser2 and eS21 on residue Met1 (**Table S3**). In addition, our MS analysis identified N-terminal acetylation of uS7, eS6, eS12, uS13 and eS19; however, N-termini of these RPs were disordered, thus impairing their direct visualization (**Table S3**). Due to the absence of confirmatory tryptic peptides, we cannot exclude the possibility that additional 40S subunit ribosomal proteins may also be acetylated. We set out to obtain insight into the structural role of the N-terminal acetylation of uS2 and eS21. The N-terminus of uS2 model is predicted with low (70 > pLDDT > 50; pLDDT: per-residue confidence score) confidence by AlphaFold (AF, (Jumper et al. 2021)), indicating it may be unstructured. Superposing the uS2 model derived from our reconstruction with the AF predicted model (RMSD over 206 residues: 0.65 Å), suggests that acetylation may promote the interaction of the Ser2 N-terminus with residue Arg63 (**Fig. 3 C**). In contrast, although the cryo-EM and AlphaFold models of eS21 superimpose well (RMSD over 83 residues: 0.48 Å), the C-terminal loop of uS5 is displaced away from the N-terminal acetylated Met1 of eS21 (**Fig. 3 D**). This analysis suggests that N-terminal acetylation of RPs may promote different local interactions within the ribosome.

**Figure 3:**
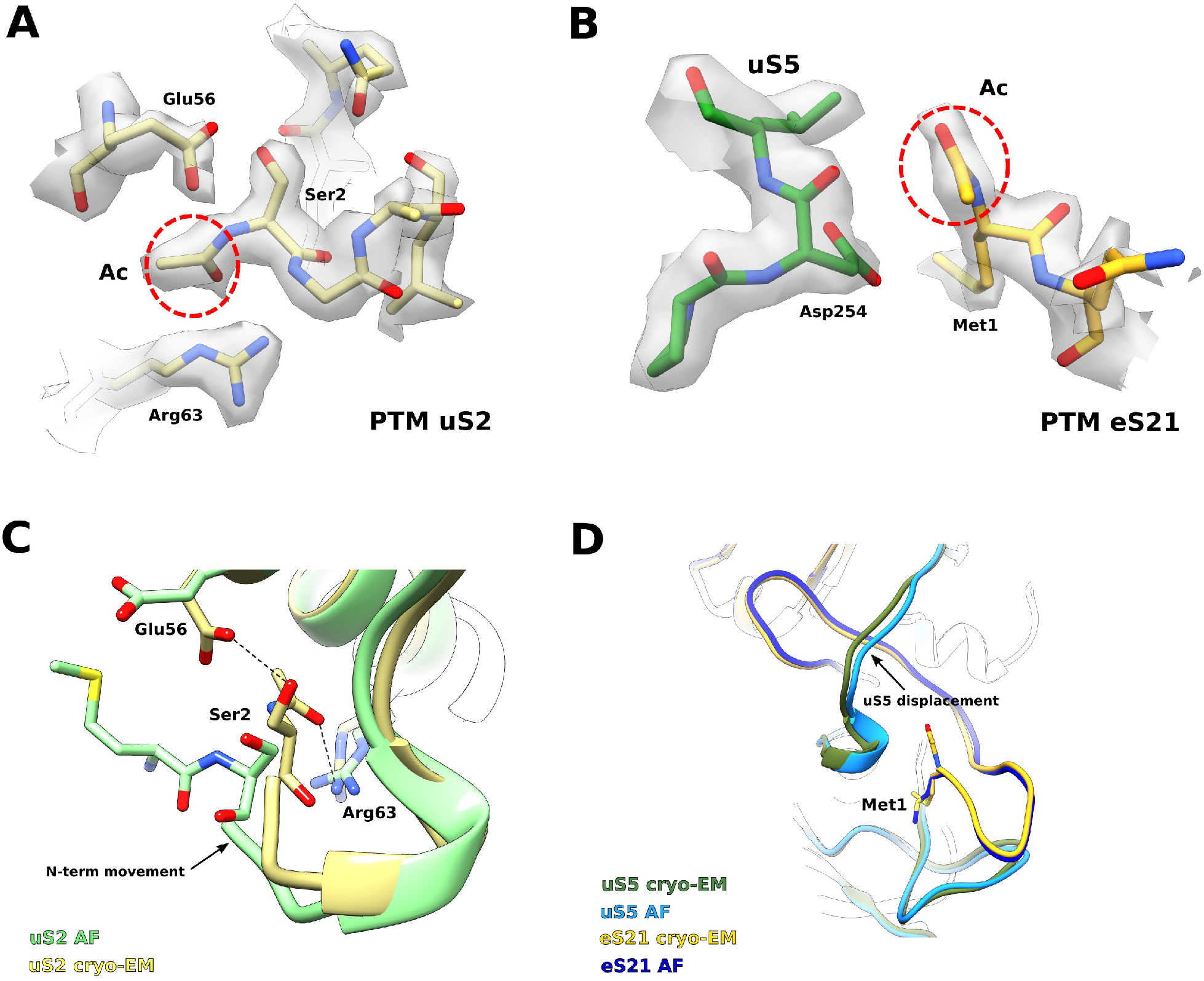
Visualizing post-translational modifications of the human 40S subunit. A) Close-up of the N-terminal region of the universally conserved ribosomal protein uS2. N-terminal acetylation of residue Ser2 is highlighted (red circle). B) N-terminus of the eukaryotic specific eS21, with the N-terminal acetylation highlighted (red circle). C) Superposition of uS2 model derived from our reconstruction, colored sand, with the model predicted by AlphaFold (Jumper et al. 2021), colored light green. The first residue at the N-terminus of both models is shown as sticks, remaining residues are shown as ribbons. AF: AlphaFold. D) Superposition of i) eS21 model derived from our reconstruction, colored yellow, on the predicted model generated by AlphaFold, colored dark blue; ii) uS5 model built from our reconstruction (forest green) overlaid on the AlphaFold predicted model colored slate blue (RMSD over 219 residues: 0.61 Å). Models are shown as ribbons and acetylated Met1 as sticks.

## DISCUSSION

The structural role of monovalent and divalent cations in ribosomal function has long remained a matter of debate. Although recent developments in long-wavelength X-ray diffraction allowed experimental assignment of K^+^ and Mg^2+^ in the bacterial ribosome (Rozov et al. 2019), the high resolution that we have achieved in our reconstructions has allowed us to identify the position of ions directly and unambiguously in the most stable regions of the human 40S ribosomal subunit (**Fig. 2**). Consistent with a role for Mg^2+^ ions in stabilizing and promoting rRNA folding, we visualized about 90% of them bound to the phosphate backbone of the 18S with octahedral coordination. K^+^ ions instead bind to different elements of the ribosome to: i) stabilize the interaction between a RP and rRNA via square anti-prismatic coordination (**Fig. 2 A, C, E**) or ii) promote rRNA nucleobase stacking (**Fig. 2 B, D, G**). In several regions of the 40S ribosomal subunit, we observed K^+^ ions bound in identical positions compared to the bacterial ribosome, promoting similar bonding patterns (**Fig. 2 A-D and S5 A-D**). In addition, we identified K^+^ ions bound in eukaryote-specific positions (**Fig. 2 E-G**), mediating similar structural roles.

RNA modifications are an important source of ribosome heterogeneity that may potentially modulate ribosome function and gene expression during normal development and in cancer (Caizergues-Ferrer et al. 1990; Hebras et al. 2020; Bouffard et al. 2018; Delhermite et al. 2021; Marcel et al. 2013; Su et al. 2014; McMahon et al. 2015; Liang et al. 2019; Barros-Silva et al. 2021; Zhang et al. 2019; Monaco et al. 2018; Janin et al. 2020). The human 40S ribosomal subunit alone contains 10 of the 14 identified rRNA post-transcriptional modifications (Taoka et al. 2018), which are mainly found in functional regions such as the mRNA path, the decoding center (DC) and the binding platforms for translation and assembly factors, as well as the core of the head and body domains, where they stabilize rRNA structure. Our reconstructions support the hypothesis that chemical modifications of rRNA fulfill structural and regulatory roles during ribosome biogenesis and translation. Of particular interest is the set of rRNA modifications within the h8-h14 region, where three 2’-O-methylations (C_m_462 and A_m_468, directly on h14; A_m_159, tip of h8) are incorporated (**Fig. S6 A**). These modifications all lie within van der Waals radius distance, which may help promote h14 folding. In bacteria, mutations in residues within h8-h14 promote miscoding *in vivo* (McClory et al. 2010), attesting to the functional importance of this region for translation. In eukaryotes, h8-h14 forms the binding platform for several translation factors, such as the ATPase ABCE1 (involved in recycling) and the GTPases eIF5B and eIF1α (initiation), eEF2 (elongation) and eRF3 (termination). Due to its proximity to the nucleotide-binding domain of translation factors, the h8-h14 region may trigger nucleotide hydrolysis by these factors (des Georges et al. 2014; Preis et al. 2014; Shao et al. 2016) (**Fig. S6 B**). However, the underlying molecular mechanism remains elusive. In eukaryotes, the neck-head junction at h29 contains a conserved guanosine residue at position 1639, which carries an N^7^-methylation (**Fig. S3 and S6 C-D**). During translocation, h29 participates in movement of the tRNA from the P- to the E-site, with residues G1639 and the neighboring A1640 directly interacting with its anticodon stem loop (**Fig. S6 D**).

Our cryo-EM reconstructions have allowed us to visualize post-transcriptional modifications in unprecedented detail (**Fig. 1, Fig. S3 and S4**). Achieving high-resolution maps of the 40S ribosomal subunit is a challenging task, given the presence of rRNA elements that are highly dynamic in absence of the large ribosomal subunit. In particular, h44, which constitutes the DC, spans the entire length of the body and a large portion of it is flexible when not interacting with H69-71 of the 60S ribosomal subunit. Nonetheless, our reconstruction allowed to visualize most of the post-transcriptional modifications incorporated within the 18S rRNA, including the unique rRNA modifications at the base of h44 (**Fig. S4**). However, we were only able to visualize partial density for the hypermodified 1-methyl-3-(3-amino-3-carboxypropyl) ψ1248, confirming its high mobility when it is not bound to initiator tRNA (Simonetti et al. 2020; Brito Querido et al. 2020) or to the assembly factor RIOK1 (Ameismeier et al. 2020). Consistent with the SILNAS-based analysis of human 80S ribosomes (Taoka et al. 2018), we visualized the previously unmodeled human rRNA modifications A_m_590, G_m_1447, ψ572, ψ863 and ψ1136 (**Fig. 1, Fig. S3 and S4**). Surprisingly, we were unable to validate one of the modifications recently identified by SILNAS-based quantification analysis, 2’-O-methylation of rRNA residue C621 (Taoka et al. 2018), observing by contrast that the 2’-hydroxyl is bound to a structured water molecule (**Fig. S4 A**). We speculate that this discrepancy may relate to the different cells of origin for the 40S subunit samples (HEK293, this study, Ameismeier et al. 2020; TK6 and HeLa cells, Taoka et al. 2018). However, more experimental evidence will be required to further test this hypothesis.

The pipeline we have established enables cryo-EM reconstructions of human 40S subunits at resolutions close to 2 Å, allowing us to begin to unveil the structural basis of ribosome specialization. Additionally, we anticipate that this work will pave the way for high-resolution studies of human translation initiation complexes to explore the role of rRNA modifications and ion composition on ribosome function.

## MATERIALS AND METHODS

### Purification of human 40S ribosomal subunits

Human Embryonic Kidney Expi293 cells were grown to a density of approximately 2×10^6^ in Expi293 medium (Thermo Scientific). Following stress induction upon addition of 100 mM (final concentration) NaCl to 250 mL of culture for 20 min, cells were harvested by centrifugation at 800 x g for 10 min. After two consecutive washes with phosphate buffer saline (PBS), cells were harvested by centrifugation, flash frozen in liquid N_2_ and stored at −80 °C. Cells were thawed on ice for 30 min and later resuspended in lysis buffer (30 mM Hepes-KOH pH 7.5, 50 mM KCl, 10 mM MgCl_2_, 220 mM sucrose, 2 mM DTT, 0.5 mM EDTA, supplemented with 0.5 mM NaF, 0.1 mM Na_3_VO_4_, 0.5 x protein inhibitors cocktail tablet (Roche), 2000 units of RNasin Plus ribonuclease inhibitor (Promega), 0.5 mg/mL heparin and 0.5% Igepal-630 (Sigma)). The slurry was incubated for 30 min on ice with periodic mixing, then centrifuged at 10,000 x g for 5 min at 4 °C. The recovered soluble fraction was centrifuged at 30,000 x g for 20 min at 4 °C. The resulting supernatant was placed in a separate tube, incubated on ice for 15 min and following the addition of PEG 20,000 to a final concentration of 1.35 %, centrifuged at 20,000 x g for 12 min at 4 °C. The supernatant was placed in a clean tube and the KCl raised to a final concentration of 130 mM. The solution was incubated for 5 min and cleared by centrifugation at 17,500 x g for 5 min. The concentration of PEG 20,000 was raised to 4.25% and the solution incubated for 10 min in ice. After centrifugation at 17,500 x g for 10 min at 4 °C the white ribosome-containing pellet was resuspended in 750 μL of buffer R (30 mM Hepes-KOH pH 7.5, 125 mM KCl, 7.5 mM MgCl_2_, 2 mM DTT containing 0.5 mg/mL heparin and 0.1 x protease inhibitor cocktail) on ice. 80 OD_260_ were loaded per 10-30% sucrose gradient, prepared in buffer containing 25 mM Hepes-KOH pH 7.5, 125 mM KCl, 8.3 mM MgCl_2_, 0.5 mM EDTA, 2 mM DTT. Gradients were centrifuged at 42,800 x g for 15 h at 4 °C in an SW28 rotor (Beckman Coulter). Fractions of 1 mL were collected using an AKTA system (Cytiva). Those fractions corresponding to the 80S peak were pooled and precipitated by adding PEG 20,000 at a final concentration of 5.15%. The solution was incubated for 10 min at 4 °C and centrifuged at 17,500 x g for 10 min at 4 °C. The resulting pellet was resuspended in buffer S (25 mM Hepes-KOH pH 7.5, 100 mM KCl, 7.5 mM MgCl_2_ and 2 mM DTT). 70 OD_260_ were layered onto a 10-40% sucrose gradient prepared in buffer containing 25 mM Hepes pH 7.5, 550 mM KCl, 5 mM MgCl _2_ and 2 mM DTT and centrifuged at 70,000 x g for 20.5 h at 4 °C in an SW28 rotor (Beckman Coulter). Fractions corresponding to the 40S peak were collected, pooled and 40S subunits precipitated by adding 7.5% of PEG 20,000 (final concentration) for 10 min on ice and centrifuging at 17,500 x g for 10 min at 4 °C. The final pellet was resuspended in buffer containing 20 mM Hepes-KOH pH7.5, 100 mM KCl, 5 mM MgOAc2 and 1 mM DTT. Aliquots (final concentration of 35 OD_260_/μL) were flash frozen in liquid N2 and stored at −80°C.

### Cryo-EM grids preparation, data collection and image processing

R2/2 holey carbon supported copper grids (Quantifoil) coated with a 2nm thin carbon layer were treated as described (A. Faille et al., in preparation). 4 μL of 40S sample (at a concentration of 2.5 OD_260_/μL) was directly applied onto each grid. The grid was incubated for 30 sec at 4 °C and 95% humidity, then blotted for 1 s (blot force −7) prior to plunging into liquid ethane using a Vitrobot Mark IV (FEI Company). Grids were screened for ice quality and cryo-EM data was acquired on a Titan Krios transmission electron microscope (FEI Company) at 300 kV equipped with a K3 direct electron detector (Gatan). The dataset was recorded in counting super-resolution mode at a nominal magnification of 105,000x (corresponding to a pixel size of 0.83 Å/px on the object scale (0.415 Å in super-resolution) with a defocus range of −1.0 to −2.8 μm and a dose of ~49 e^−^/Å^2^. Acquisition of 3091 movies was performed semi-automatically using EPU software (FEI Company) and pre-processed on-the-fly using Warp (Tegunov and Cramer 2019). The resulting set of picked particles was imported into cryoSPARC (Punjani et al. 2017) to perform initial processing and remove non-ribosomal and/or damaged particles from the dataset. After *ab initio* reconstruction, heterogeneous and uniform refinement, a total of 474276 particles were imported into Relion 3.0 (Zivanov et al. 2018) for further processing. Original movies were corrected for the effects of beam-induced motion using MotionCor2 (Zheng et al. 2017) as implemented in the Relion GUI, and particles from cryoSPARC re-extracted. A first round of Refine3D was performed using the map resulting from homogeneous refinement in cryoSPARC as a reference. The estimated resolution after postprocessing was 2.41 Å. CTF and beam tilt refinement followed by Bayesian polishing was repeated twice, and Refine3D performed after each step. Final postprocessing of the entire human 40S yielded a reconstruction at 2.15 Å resolution, with a B-factor of −35 Å^2^. The alignment of the particles is driven by the body domain of the 40S subunit, as the head is mobile. To achieve high-resolution reconstructions for both domains, two overlapping soft masks including the individual domains were created within Relion and used as inputs for multibody refinement (Nakane and Scheres 2021). The two resulting reconstructions yielded a resolution estimation of 2.24 Å (B-factor: −39 Å^2^) for the head domain and 2.09 Å (B-factor: −33 Å^2^) for the body domain. All resolution estimations are based on the gold-standard Fourier Shell Correlation (FSC) criterion of 0.143 (Henderson et al. 2012; Scheres and Chen 2012).

### Model building and validation

The resulting maps were used for model building in Coot (Emsley and Cowtan 2004) using the mature 40S ribosomal subunit (PDB: 6G5H) as a reference. Protein and rRNA chains were inspected in Coot and manually fitted into the density. To interpret regions at lower local resolution, we used LocalDeblur (Ramírez-Aportela et al. 2020) from the Scipion suite (de la Rosa-Trevín et al. 2016) to generate locally sharpened maps of the head and the body domains. To add water molecules, we employed a semi-automatic approach using phenix.douse (Phenix suite, (Liebschner et al. 2019)) and visual inspection in Coot. Building of additional water molecules and ions, such as Mg^2+^ and K^+^, was performed manually in Coot, based on the coordination geometries. Models were refined using real space refinement as implemented in PHENIX (Liebschner et al. 2019) and validated using MolProbity (Williams et al. 2018).

### Mass spectrometry

A sample containing 30 OD_260_ of the human 40S subunit was loaded onto a 4-12% acrylamide gel and electrophoresed for 5 min to remove excess sucrose. Gel bands were excised, destained, reduced (using DTT), alkylated (iodoacetamide) and subjected to enzymatic digestion with trypsin (Promega) overnight at 37 °C. After digestion, the supernatant was pipetted into a sample vial and loaded onto an autosampler for automated LC-MS/MS analysis. The liquid chromatography (LC)-MS/MS experiment was performed using a Dionex Ultimate 3000 RSLC nanoUPLC (Thermo Fisher Scientific) system and a Q Exactive Orbitrap mass spectrometer (Thermo Fisher Scientific). Separation of peptides was performed by reverse-phase chromatography at a flow rate of 300 nL/min and a Thermo Scientific reverse-phase nano Easy-spray column (Thermo Scientific PepMap C18, 2 μm particle size, 100 Å pore size, 75 μm i.d. x 50 cm length). Peptides were loaded onto a pre-column (Thermo Scientific PepMap 100 C18, 5 μm particle size, 100 Å pore size, 300 μm i.d. x 5 mm length) from the Ultimate 3000 autosampler with 0.1 % formic acid for 3 min at a flow rate of 15 μL/min. After this period, the column valve was switched to allow elution of peptides from the pre-column onto the analytical column. Solvent A was water + 0.1 % formic acid and solvent B was 80 % acetonitrile, 20 % water + 0.1 % formic acid. The linear gradient employed was 2-40 % B over 90 min (the total run time including column washing and re-equilibration was 120 min). The LC eluant was sprayed into the mass spectrometer by means of an Easy-spray source (Thermo Fisher Scientific Inc.). All *m/z* values of eluting ions were measured in an Orbitrap mass analyzer, set at a resolution of 35000 and scanned between *m/z* 380-1500. Data dependent scans (Top 20) were employed to automatically isolate and generate fragment ions by higher energy collisional dissociation (HCD). Normalized collision energy (NCE): 25%) in the HCD collision cell and measurement of the resulting fragment ions was performed in the Orbitrap analyzer, set at a resolution of 17500. Singly charged ions and ions with unassigned charge states were excluded from selection for MS/MS and a dynamic exclusion of 60 seconds was employed.

The files were submitted to the Mascot search algorithm (Matrix Science, London UK, version 2.6.0) and searched against a common contaminants database (125 sequences; 41129 residues) and the UniProt human database (CCP_UniProt_homo sapiens_proteome_20180409 database, 93609 sequences; 37041084 residues). Variable modifications of oxidation (M), deamidation (NQ) phosphorylation (S,T and Y) acetylation (K and protein N-terminus), methylation (HKNR), sulfation (STY) and ubiquitination (K) were applied together with a fixed modification of carbamidomethyl (C). The peptide and fragment mass tolerances were set to 20 ppm and 0.1 Da, respectively. A significance threshold value of p<0.05 and a peptide cut-off score of 20 were also applied. Scaffold (version Scaffold_4.10.0, Proteome Software Inc., Portland, OR) was used to validate MS/MS based peptide and protein identifications. Peptide identifications were accepted if they could be established at greater than 95.0 % probability by the Peptide Prophet algorithm (Keller et al. 2002) with Scaffold delta-mass correction. Protein identifications were accepted if they could be established at greater than 99.0 % probability and contained at least 2 identified peptides.

Protein probabilities were assigned by the Protein Prophet algorithm (Nesvizhskii et al. 2003). Proteins that contained similar peptides and could not be differentiated based on MS/MS analysis alone were grouped to satisfy the principles of parsimony. Proteins sharing significant peptide evidence were grouped into clusters.

## DATA DEPOSITON

Coordinates and cryo-EM density maps were deposited in the RCSB Protein Data Bank with accession code PDB-xxxx for the 40S ribosomal subunit model and the Electron Microscopy Database with accession codes EMD-xxxx, EMD-xxxx and EMD-xxxx for the entire human 40S, head and body domains, respectively.

## ACKNOWLEDGMENTS

Proteomics experiments were performed at the Cambridge Centre for Proteomics. We thank the laboratory of Dr. Venki Ramakrishnan for providing Expi293 cells.

## SUPPLEMENTARY FIGURE LEGENDS

**Figure S1:**
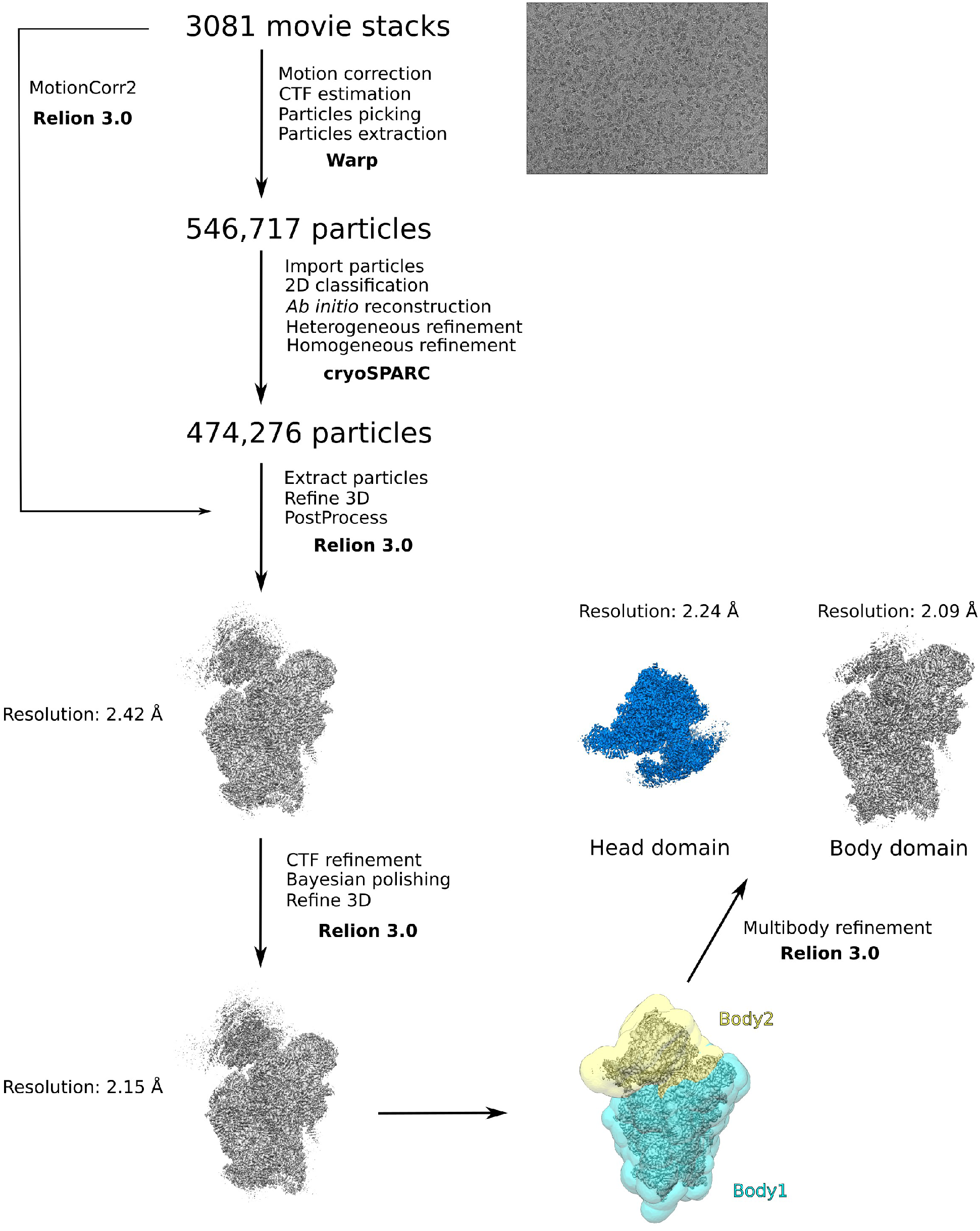
Cryo-EM data processing scheme.

**Figure S2:**
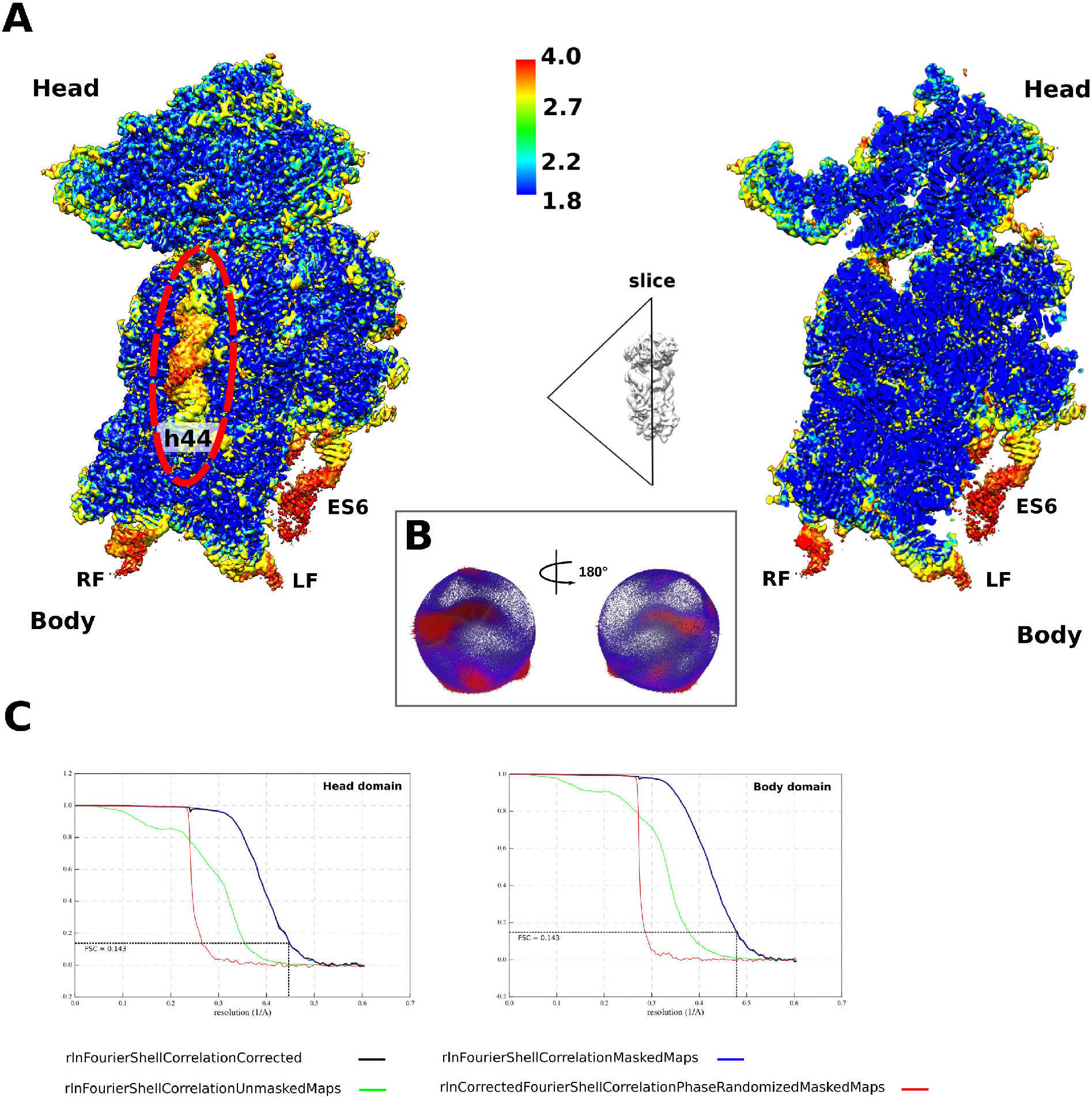
Local resolution of human 40S subunit extends to 1.8 Å. A) Top left: local resolution estimate of the complete human 40S subunit, viewed from the intersubunit interface. The less well-resolved regions are indicated: h44 (red dashed line), right (RF) and left (LF) feet, rRNA expansion segment 6 (ES6). Top right: slice through the 40S showing high-resolution within the most stable regions of the 40S ribosomal subunit body and head domains. The color scale indicates the resolution estimation values (Angstrom). B) Graphical representation of particle orientation distribution during 3D refinement (over-represented views in red). C) FSC curves, as calculated within Relion 3.0 (Zivanov et al. 2018), for the head and the body domain. Gold-standard FSC cut-off at 0.143 to estimate the final resolution, is shown.

**Figure S3:**
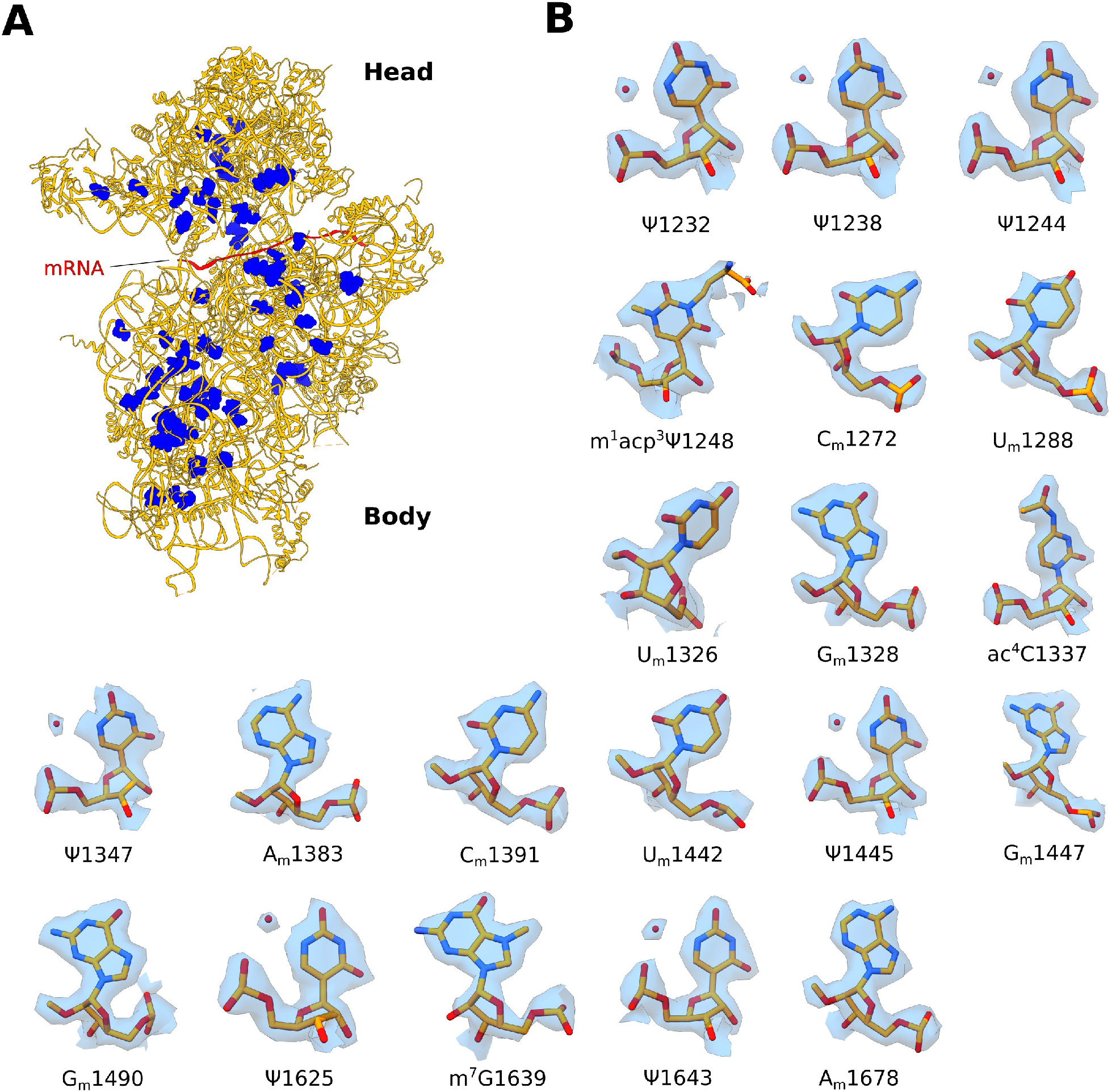
rRNA modifications of the head domain. A) Ribbon representation of the human 40S subunit (orange) with rRNA modifications shown as blue spheres. For clarity, mRNA was modeled *in silico* by superposing the human 48S initiation complex (PDB: 6ZMW). B) rRNA modifications visualized in this study fitted into the experimental cryo-EM map, for the head domain only. Water molecules (red spheres) coordinated with pseudouridine bases are indicated.

**Figure S4:**
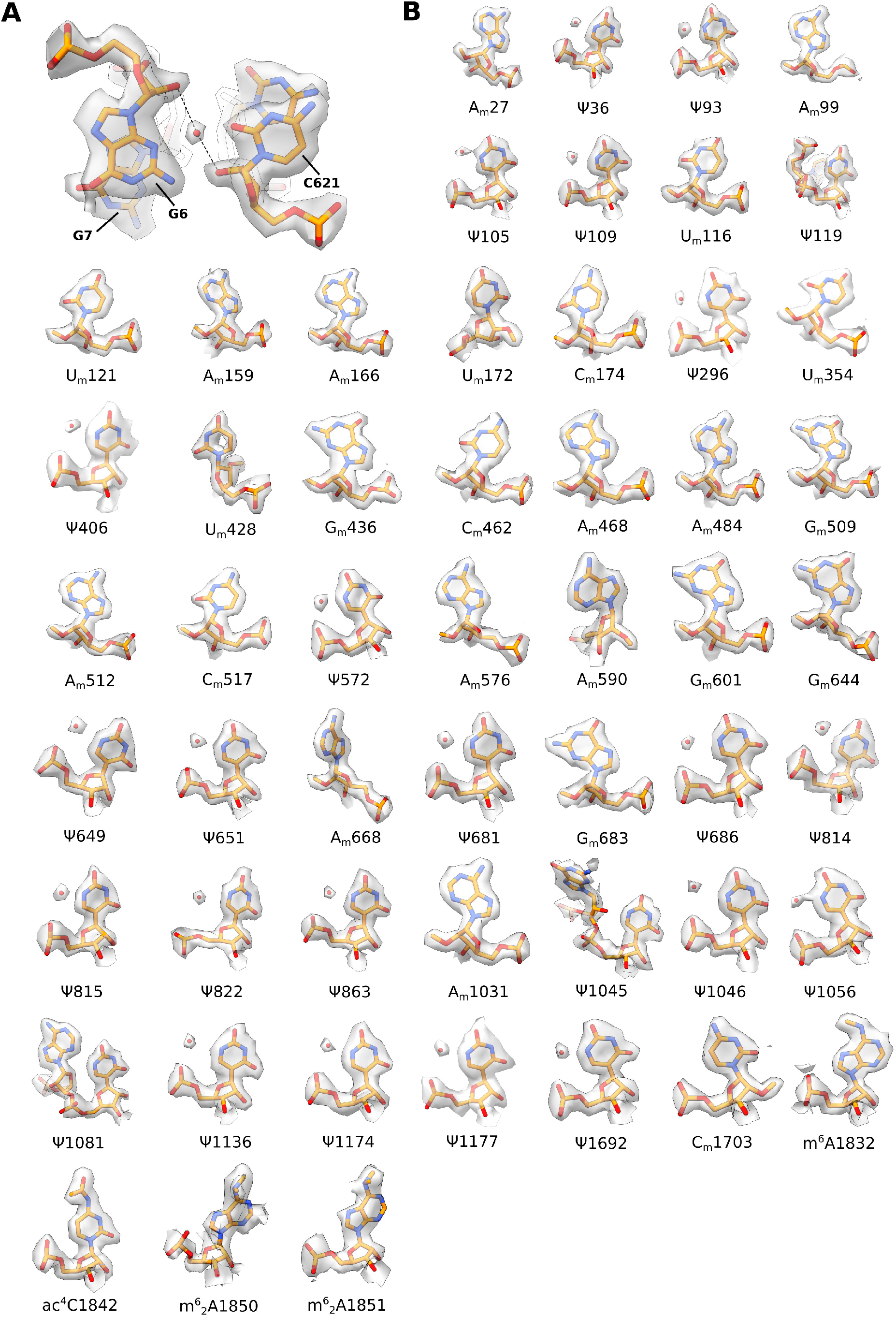
rRNA modifications in the body domain of the human 40S ribosomal subunit. A) Detail of the universally conserved rRNA residue C621 (C525 bacterial numbering) on h18. A water molecule (red sphere) coordinated with the 2’-O of the sugar backbone of C621 is conserved between higher eukaryotes and prokaryotes (PDB: 7K00). B) rRNA modifications within the body. Water molecules (red spheres) coordinated with pseudouridine bases are shown.

**Figure S5:**
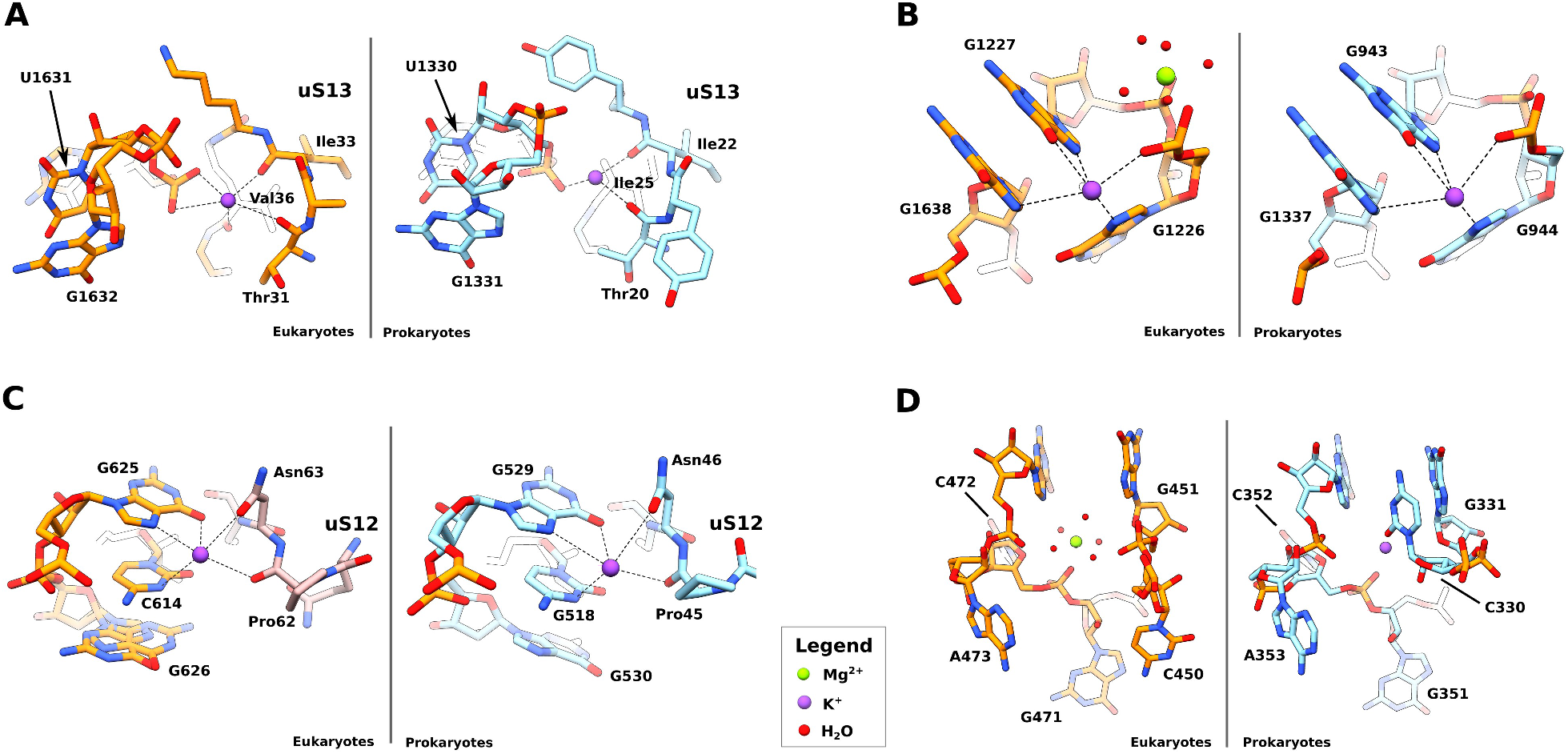
Conserved 40S ribosomal subunit solvation. Comparison between human vs *Thermus thermophilus* small ribosomal subunit showing: A) K^+^ ion stabilizing the interaction between U1631 (bacterial U1330) on h42 with the universally conserved protein uS13; B) conserved potassium ion bound close to h29; C) close-up of the decoding center showing a K^+^ ion coordinating the interaction of h18 with the universally conserved ribosomal protein uS12; D) K^+^ ion promoting nucleobases stacking on h28; E) eukaryote-specific binding of Mg^2+^ to the h13-14 region. *Thermus thermophilus* PDB 6QNR was used for comparison. Water molecules, K^+^ and Mg^2+^ ions are shown as spheres (legend, bottom of panel).

**Figure S6:**
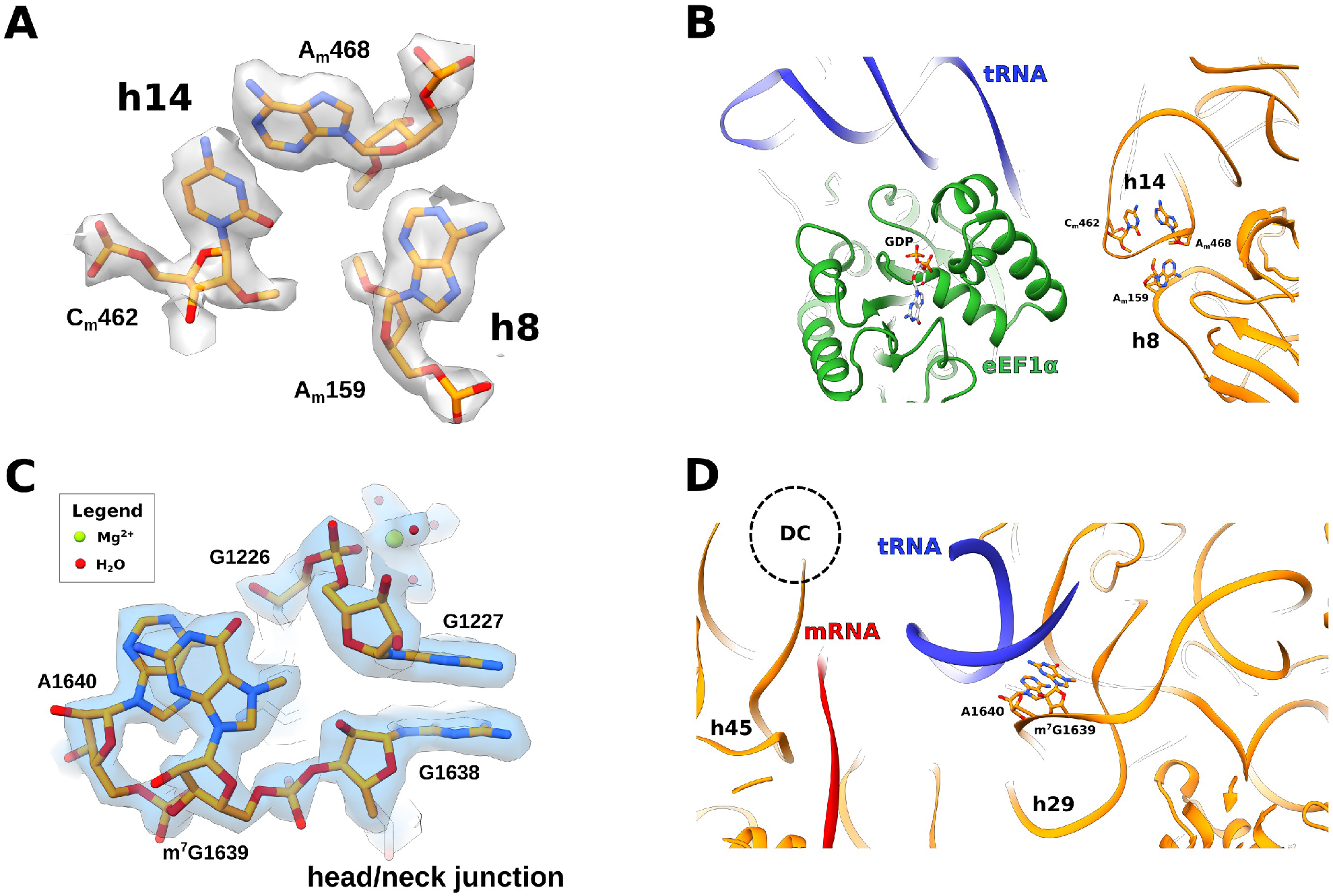
rRNA modifications essential for translation. A) Fit of rRNA residues A_m_159 (h8) A_m_428 and C_m_462 (h14) into the cryo-EM density map of the 40S body. B) Close-up view of h8-h14 interaction with the eEF1α-tRNA complex (eEF1α in green, tRNA in blue). 18S rRNA residues A_m_159, C_m_462 and A_m_468 and the GDP molecule within eEF1α are shown as sticks. C) Detail of the head/neck junction region on the head, with a close-up view of m^7^G1639 and neighboring rRNA residues, fitted within the cryo-EM density of the head. Water molecules and Mg^2+^ ions are shown as spheres (legend, top left). D) Interaction between rRNA residue m^7^G1639 and P-site tRNA (in blue), upon superposition with human 48S initiation complex (PDB: 6ZMW). The mRNA (red) is shown for clarity.

**Supplementary Table S1:**

|  | Human 40S<br>(EMD-xxxx)<br>(PDB xxxx) | Head domain<br>(EMD-xxxx) | Body domain<br>(EMD-xxxx) |
| --- | --- | --- | --- |
| **Data collection and processing** |  |  |  |
| Magnification | 105,000x | 105,000x | 105,000x |
| Voltage (kV) | 300 | 300 | 300 |
| Electron exposure: total dose (e-/Å<sup>2</sup>) | 40.56 | 40.56 | 40.56 |
| Defocus range (μm) | -2.8, -2.5, -2.2, -1.9,<br>-1.6, -1.3, -1.0 | -2.8, -2.5, -2.2, -1.9,<br>-1.6, -1.3, -1.0 | -2.8, -2.5, -2.2, -1.9,<br>-1.6, -1.3, -1.0 |
| Pixel size (Å) | 0.83 (0.415 super-res) | 0.83 (0.415 super-res) | 0.83 (0.415 super-res) |
| Symmetry imposed | C1 | C1 | C1 |
| Initial particle images (no.) | 546,717 | 546,717 | 546,717 |
| Final particle images (no.) | 474,276 | 474,276 | 474,276 |
| Map resolution (Å) |  |  |  |
| FSC threshold | 2.15 | 2.24 | 2.09 |
| Map resolution range (Å) | 4.00 – 1.80 | 4.00 – 1.80 | 4.00 – 1.80 |
| Map sharpening B factor (Å<sup>2</sup>) | -35.47 | -38.64 | -33.45 |
| **Refinement** |  |  |  |
| Initial model used (PDB code) | 6G5H |  |  |
| Model resolution range (Å) | 5.00 – 2.15 |  |  |
| Model composition |  |  |  |
| Non-hydrogen atoms | 75673 |  |  |
| Protein residues | 4809 |  |  |
| RNA bases | 1656 |  |  |
| Ligands |  |  |  |
| Water | 1682 |  |  |
| Mg<sup>2+</sup> | 83 |  |  |
| K<sup>+</sup> | 81 |  |  |
| B factors (mean, Å<sup>2</sup>) |  |  |  |
| Protein | 26.98 |  |  |
| RNA | 28.89 |  |  |
| Ligand | 23.68 |  |  |
| Water | 19.48 |  |  |
| R.m.s. deviations |  |  |  |
| Bond lengths (Å) | 0.006 |  |  |
| Bond angles (°) | 0.785 |  |  |
| Validation |  |  |  |
| MolProbity score (percentile, 0-99<br>Å) | 1.91 (80<sup>th</sup>)<br>5.12 (93<sup>rd</sup>) |  |  |
| Clashscore (all atoms) | 3.59 |  |  |
| Poor rotamers (%) |  |  |  |
| Ramachandran plot |  |  |  |
| Favored (%) | 96.64 |  |  |
| Allowed (%) | 3.30 |  |  |
| Disallowed (%) | 0.06 |  |  |
| CaBLAM outliers (%) | 2.00 |  |  |
| **Model vs. Map validation** |  |  |  |
| CC<sub>mask</sub> | 0.84 |  |  |
| CC<sub>box</sub> | 0.79 |  |  |
| CC<sub>volume</sub> | 0.81 |  |  |
| FSC model-map (0.5) (Å) | 2.2 |  |  |
| **RNA** |  |  |  |
| Correct sugar packers (%) | 99.70 |  |  |
| Good backbone conformations (%) | 83.95 |  |  |

**Supplementary Table S2:**
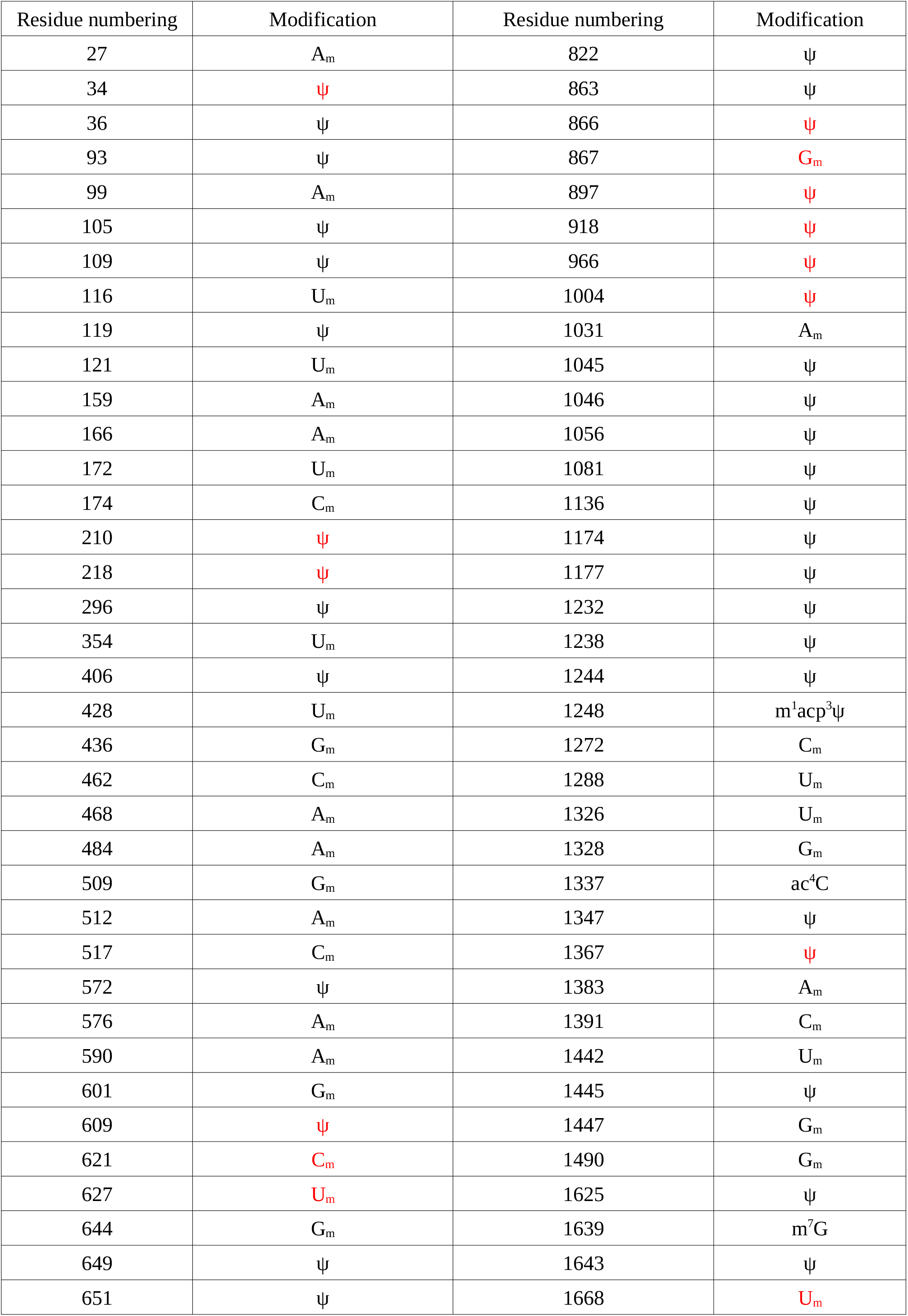
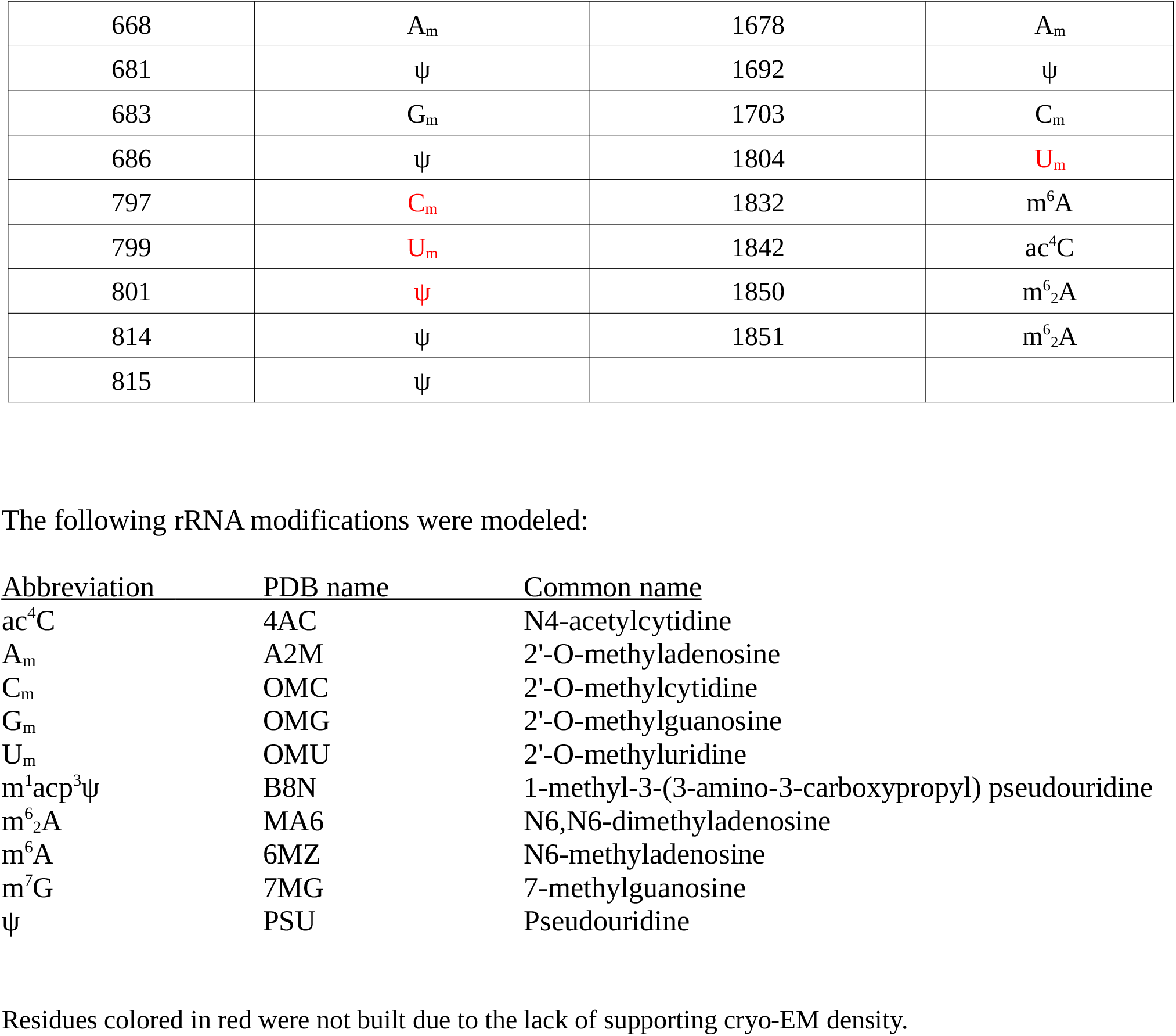

| Residue numbering | Modification | Residue numbering | Modification |
| --- | --- | --- | --- |
| 27 | A<sub>m</sub> | 822 | ψ |
| 34 | ψ | 863 | ψ |
| 36 | ψ | 866 | ψ |
| 93 | ψ | 867 | G<sub>m</sub> |
| 99 | A<sub>m</sub> | 897 | ψ |
| 105 | ψ | 918 | ψ |
| 109 | ψ | 966 | ψ |
| 116 | U<sub>m</sub> | 1004 | ψ |
| 119 | ψ | 1031 | A<sub>m</sub> |
| 121 | U<sub>m</sub> | 1045 | ψ |
| 159 | A<sub>m</sub> | 1046 | ψ |
| 166 | A<sub>m</sub> | 1056 | ψ |
| 172 | U<sub>m</sub> | 1081 | ψ |
| 174 | C<sub>m</sub> | 1136 | ψ |
| 210 | ψ | 1174 | ψ |
| 218 | ψ | 1177 | ψ |
| 296 | ψ | 1232 | ψ |
| 354 | U<sub>m</sub> | 1238 | ψ |
| 406 | ψ | 1244 | ψ |
| 428 | U<sub>m</sub> | 1248 | m<sup>1</sup>acp<sup>3</sup>ψ |
| 436 | G<sub>m</sub> | 1272 | C<sub>m</sub> |
| 462 | C<sub>m</sub> | 1288 | U<sub>m</sub> |
| 468 | A<sub>m</sub> | 1326 | U<sub>m</sub> |
| 484 | A<sub>m</sub> | 1328 | G<sub>m</sub> |
| 509 | G<sub>m</sub> | 1337 | ac<sup>4</sup>C |
| 512 | A<sub>m</sub> | 1347 | ψ |
| 517 | C<sub>m</sub> | 1367 | ψ |
| 572 | ψ | 1383 | A<sub>m</sub> |
| 576 | A<sub>m</sub> | 1391 | C<sub>m</sub> |
| 590 | A<sub>m</sub> | 1442 | U<sub>m</sub> |
| 601 | G<sub>m</sub> | 1445 | ψ |
| 609 | ψ | 1447 | G<sub>m</sub> |
| 621 | C<sub>m</sub> | 1490 | G<sub>m</sub> |
| 627 | U<sub>m</sub> | 1625 | ψ |
| 644 | G<sub>m</sub> | 1639 | m<sup>7</sup>G |
| 649 | ψ | 1643 | ψ |
| 651 | ψ | 1668 | U<sub>m</sub> |
| 668 | A<sub>m</sub> | 1678 | A<sub>m</sub> |
| 681 | ψ | 1692 | ψ |
| 683 | G<sub>m</sub> | 1703 | C<sub>m</sub> |
| 686 | ψ | 1804 | U<sub>m</sub> |
| 797 | C<sub>m</sub> | 1832 | m<sup>6</sup>A |
| 799 | U<sub>m</sub> | 1842 | ac<sup>4</sup>C |
| 801 | ψ | 1850 | m<sup>6</sup><sub>2</sub>A |
| 814 | ψ | 1851 | m<sup>6</sup><sub>2</sub>A |
| 815 | ψ |  |  |

The following rRNA modifications were modeled:
| Abbreviation | PDB name | Common name |
| --- | --- | --- |
| ac<sup>4</sup>C | 4AC | N4-acetylcytidine |
| A<sub>m</sub> | A2M | 2'-O-methyladenosine |
| C<sub>m</sub> | OMC | 2'-O-methylcytidine |
| G<sub>m</sub> | OMG | 2'-O-methylguanosine |
| U<sub>m</sub> | OMU | 2'-O-methyluridine |
| m<sup>1</sup>acp<sup>3</sup>ψ | B8N | 1-methyl-3-(3-amino-3-carboxypropyl) pseudouridine |
| m<sup>6</sup><sub>2</sub>A | MA6 | N6,N6-dimethyladenosine |
| m<sup>6</sup>A | 6MZ | N6-methyladenosine |
| m<sup>7</sup>G | 7MG | 7-methylguanosine |
| ψ | PSU | Pseudouridine |

Residues colored in red were not built due to the lack of supporting cryo-EM density.

## REFERENCES

Ameismeier M, Cheng J, Berninghausen O, Beckmann R. 2018. Visualizing late states of human 40S ribosomal subunit maturation. Nature 558: 249–253. doi: 10.1038/s41586-018-0193-0

Ameismeier M, Zemp I, van den Heuvel J, Thoms M, Berninghausen O, Kutay U, Beckmann R. 2020. Structural basis for the final steps of human 40S ribosome maturation. Nature 587: 683–687. doi: 10.1038/s41586-020-2929-x

Aubert M, O’Donohue M-F, Lebaron S, Gleizes P-E. 2018. Pre-Ribosomal RNA Processing in Human Cells: From Mechanisms to Congenital Diseases. Biomolecules 8: 123. doi: 10.3390/biom8040123

Barbieri I, Kouzarides T. 2020. Role of RNA modifications in cancer. Nat Rev Cancer 20: 303–322. doi: 10.1038/s41568-020-0253-2

Barros-Silva D, Klavert J, Jenster G, Jerónimo C, Lafontaine DLJ, Martens-Uzunova ES. 2021. The role of OncoSnoRNAs and Ribosomal RNA 2’-O-methylation in Cancer. RNA Biol 18: 61–74. doi: 10.1080/15476286.2021.1991167

Bhatt PR, Scaiola A, Loughran G, Leibundgut M, Kratzel A, Meurs R, Dreos R, O’Connor KM, McMillan A, Bode JW, et al. 2021. Structural basis of ribosomal frameshifting during translation of the SARS-CoV-2 RNA genome. Science 372: 1306–1313. doi: 10.1126/science.abf3546

Bohnsack KE, Bohnsack MT. 2019. Uncovering the assembly pathway of human ribosomes and its emerging links to disease. EMBO J 38: e100278. doi: 10.15252/embj.2018100278

Bouffard S, Dambroise E, Brombin A, Lempereur S, Hatin I, Simion M, Corre R, Bourrat F, Joly J-S, Jamen F. 2018. Fibrillarin is essential for S-phase progression and neuronal differentiation in zebrafish dorsal midbrain and retina. Dev Biol 437: 1–16. doi: 10.1016/j.ydbio.2018.02.006

Brito Querido J, Sokabe M, Kraatz S, Gordiyenko Y, Skehel JM, Fraser CS, Ramakrishnan V. 2020. Structure of a human 48S translational initiation complex. Science 369: 1220–1227. doi: 10.1126/science.aba4904

Caizergues-Ferrer M, Mathieu C, Mariottini P, Amalric F, Amaldi F. 1990. Fibrillarin and U3 RNA expression during Xenopus oogenesis and embryo development. Mol Biol Rep 14: 107–108. doi: 10.1007/BF00360434

Carlile TM, Rojas-Duran MF, Zinshteyn B, Shin H, Bartoli KM, Gilbert WV. 2014. Pseudouridine profiling reveals regulated mRNA pseudouridylation in yeast and human cells. Nature 515: 143–146. doi: 10.1038/nature13802

de la Rosa-Trevín JM, Quintana A, Del Cano L, Zaldívar A, Foche I, Gutiérrez J, Gómez-Blanco J, Burguet-Castell J, Cuenca-Alba J, Abrishami V, et al. 2016. Scipion: A software framework toward integration, reproducibility and validation in 3D electron microscopy. J Struct Biol 195: 93–99. doi: 10.1016/j.jsb.2016.04.010

Delhermite J, Tafforeau L, Sharma S, Marchand V, Wacheul L, Lattuca R, Desiderio S, Motorin Y, Bellefroid E, Lafontaine DLJ. 2021. Systematic mapping of rRNA 2’-O methylation during frog development and involvement of the methyltransferase Fibrillarin in eye and craniofacial development in *Xenopus laevis*. BioRxiv 2021.11.25.469989. doi: https://doi.org/10.1101/2021.11.25.469989

des Georges A, Hashem Y, Unbehaun A, Grassucci RA, Taylor D, Hellen CUT, Pestova TV, Frank J. 2014. Structure of the mammalian ribosomal pre-termination complex associated with eRF1.eRF3.GDPNP. Nucleic Acids Res 42: 3409–3418. doi: 10.1093/nar/gkt1279

Emsley P, Cowtan K. 2004. Coot: model-building tools for molecular graphics. Acta Crystallogr D Biol Crystallogr 60: 2126–2132. doi: 10.1107/S0907444904019158

Erales J, Marchand V, Panthu B, Gillot S, Belin S, Ghayad SE, Garcia M, Laforêts F, Marcel V, Baudin-Baillieu A, et al. 2017. Evidence for rRNA 2’-O-methylation plasticity: Control of intrinsic translational capabilities of human ribosomes. Proc Natl Acad Sci U S A 114: 12934–12939. doi: 10.1073/pnas.1707674114

Gong J, Li Y, Liu C-J, Xiang Y, Li C, Ye Y, Zhang Z, Hawke DH, Park PK, Diao L, et al. 2017. A Pan-cancer Analysis of the Expression and Clinical Relevance of Small Nucleolar RNAs in Human Cancer. Cell Rep 21: 1968–1981. doi: 10.1016/j.celrep.2017.10.070

Hebras J, Krogh N, Marty V, Nielsen H, Cavaillé J. 2020. Developmental changes of rRNA ribose methylations in the mouse. RNA Biol 17: 150–164. doi: 10.1080/15476286.2019.1670598

Heiss NS, Knight SW, Vulliamy TJ, Klauck SM, Wiemann S, Mason PJ, Poustka A, Dokal I. 1998. X-linked dyskeratosis congenita is caused by mutations in a highly conserved gene with putative nucleolar functions. Nat Genet 19: 32–38. doi: 10.1038/ng0598-32

Henderson R, Sali A, Baker ML, Carragher B, Devkota B, Downing KH, Egelman EH, Feng Z, Frank J, Grigorieff N, et al. 2012. Outcome of the first electron microscopy validation task force meeting. Structure 20: 205–214. doi: 10.1016/j.str.2011.12.014

Incarnato D, Anselmi F, Morandi E, Neri F, Maldotti M, Rapelli S, Parlato C, Basile G, Oliviero S. 2017. High-throughput single-base resolution mapping of RNA 2΄-O-methylated residues. Nucleic Acids Res 45: 1433–1441. doi: 10.1093/nar/gkw810

Janin M, Coll-SanMartin L, Esteller M. 2020. Disruption of the RNA modifications that target the ribosome translation machinery in human cancer. Mol Cancer 19: 70. doi: 10.1186/s12943-020-01192-8

Jansson MD, Häfner SJ, Altinel K, Tehler D, Krogh N, Jakobsen E, Andersen JV, Andersen KL, Schoof EM, Ménard P, et al. 2021. Regulation of translation by site-specific ribosomal RNA methylation. Nat Struct Mol Biol 28: 889–899. doi: 10.1038/s41594-021-00669-4

Jumper J, Evans R, Pritzel A, Green T, Figurnov M, Ronneberger O, Tunyasuvunakool K, Bates R, Žídek A, Potapenko A, et al. 2021. Highly accurate protein structure prediction with AlphaFold. Nature 596: 583–589. doi: 10.1038/s41586-021-03819-2

Keller A, Nesvizhskii AI, Kolker E, Aebersold R. 2002. Empirical statistical model to estimate the accuracy of peptide identifications made by MS/MS and database search. Anal Chem 74: 5383–5392. doi: 10.1021/ac025747h

Klein DJ, Moore PB, Steitz TA. 2004. The contribution of metal ions to the structural stability of the large ribosomal subunit. RNA 10: 1366–1379. doi: 10.1261/rna.7390804

Krogh N, Jansson MD, Häfner SJ, Tehler D, Birkedal U, Christensen-Dalsgaard M, Lund AH, Nielsen H. 2016. Profiling of 2’-O-Me in human rRNA reveals a subset of fractionally modified positions and provides evidence for ribosome heterogeneity. Nucleic Acids Res 44: 7884–7895. doi: 10.1093/nar/gkw482

Liang J, Wen J, Huang Z, Chen X-P, Zhang B-X, Chu L. 2019. Small Nucleolar RNAs: Insight Into Their Function in Cancer. Front Oncol 9: 587. doi: 10.3389/fonc.2019.00587

Liebschner D, Afonine PV, Baker ML, Bunkóczi G, Chen VB, Croll TI, Hintze B, Hung LW, Jain S, McCoy AJ, et al. 2019. Macromolecular structure determination using X-rays, neutrons and electrons: recent developments in Phenix. Acta Crystallogr Sect Struct Biol 75: 861–877. doi: 10.1107/S2059798319011471

Linnemann J, Pöll G, Jakob S, Ferreira-Cerca S, Griesenbeck J, Tschochner H, Milkereit P. 2019. Impact of two neighbouring ribosomal protein clusters on biogenesis factor binding and assembly of yeast late small ribosomal subunit precursors. PloS One 14: e0203415. doi: 10.1371/journal.pone.0203415

Maden BE. 1986. Identification of the locations of the methyl groups in 18 S ribosomal RNA from Xenopus laevis and man. J Mol Biol 189: 681–699. doi: 10.1016/0022-2836(86)90498-5

Maden BE. 1988. Locations of methyl groups in 28 S rRNA of Xenopus laevis and man. Clustering in the conserved core of molecule. J Mol Biol 201: 289–314. doi: 10.1016/0022-2836(88)90139-8

Maden BE, Corbett ME, Heeney PA, Pugh K, Ajuh PM. 1995. Classical and novel approaches to the detection and localization of the numerous modified nucleotides in eukaryotic ribosomal RNA. Biochimie 77: 22–29. doi: 10.1016/0300-9084(96)88100-4

Maden EH, Wakeman JA. 1988. Pseudouridine distribution in mammalian 18 S ribosomal RNA. A major cluster in the central region of the molecule. Biochem J 249: 459–464. doi: 10.1042/bj2490459

Marcel V, Ghayad SE, Belin S, Therizols G, Morel A-P, Solano-Gonzàlez E, Vendrell JA, Hacot S, Mertani HC, Albaret MA, et al. 2013. p53 acts as a safeguard of translational control by regulating fibrillarin and rRNA methylation in cancer. Cancer Cell 24: 318–330. doi: 10.1016/j.ccr.2013.08.013

McClory SP, Leisring JM, Qin D, Fredrick K. 2010. Missense suppressor mutations in 16S rRNA reveal the importance of helices h8 and h14 in aminoacyl-tRNA selection. RNA 16: 1925–1934. doi: 10.1261/rna.2228510

McMahon M, Contreras A, Ruggero D. 2015. Small RNAs with big implications: new insights into H/ACA snoRNA function and their role in human disease. Wiley Interdiscip Rev RNA 6: 173–189. doi: 10.1002/wrna.1266

Melnikov S, Ben-Shem A, Garreau de Loubresse N, Jenner L, Yusupova G, Yusupov M. 2012. One core, two shells: bacterial and eukaryotic ribosomes. Nat Struct Mol Biol 19: 560–567. doi: 10.1038/nsmb.2313

Monaco PL, Marcel V, Diaz J-J, Catez F. 2018. 2’-O-Methylation of Ribosomal RNA: Towards an Epitranscriptomic Control of Translation? Biomolecules 8: 106. doi: 10.3390/biom8040106

Nakane T, Scheres SHW. 2021. Multi-body Refinement of Cryo-EM Images in RELION. Methods Mol Biol Clifton NJ 2215: 145–160. doi: 10.1007/978-1-0716-0966-8_7

Natchiar SK, Myasnikov AG, Kratzat H, Hazemann I, Klaholz BP. 2017. Visualization of chemical modifications in the human 80S ribosome structure. Nature 551: 472–477. doi: 10.1038/nature24482

Nesvizhskii AI, Keller A, Kolker E, Aebersold R. 2003. A statistical model for identifying proteins by tandem mass spectrometry. Anal Chem 75: 4646–4658. doi: 10.1021/ac0341261

Ogle JM, Brodersen DE, Clemons WMJ, Tarry MJ, Carter AP, Ramakrishnan V. 2001. Recognition of cognate transfer RNA by the 30S ribosomal subunit. Science 292: 897–902. doi: 10.1126/science.1060612

Ojha S, Malla S, Lyons SM. 2020. snoRNPs: Functions in Ribosome Biogenesis. Biomolecules 10. doi: 10.3390/biom10050783

Pauli C, Liu Y, Rohde C, Cui C, Fijalkowska D, Gerloff D, Walter C, Krijgsveld J, Dugas M, Edemir B, et al. 2020. Site-specific methylation of 18S ribosomal RNA by SNORD42A is required for acute myeloid leukemia cell proliferation. Blood 135: 2059–2070. doi: 10.1182/blood.2019004121

Preis A, Heuer A, Barrio-Garcia C, Hauser A, Eyler DE, Berninghausen O, Green R, Becker T, Beckmann R. 2014. Cryoelectron microscopic structures of eukaryotic translation termination complexes containing eRF1-eRF3 or eRF1-ABCE1. Cell Rep 8: 59–65. doi: 10.1038/nmeth.4169

Punjani A, Rubinstein JL, Fleet DJ, Brubaker MA. 2017. cryoSPARC: algorithms for rapid unsupervised cryo-EM structure determination. Nat Methods 14: 290–296. doi: 10.1038/nmeth.4169

Ramírez-Aportela E, Vilas JL, Glukhova A, Melero R, Conesa P, Martínez M, Maluenda D, Mota J, Jiménez A, Vargas J, et al. 2020. Automatic local resolution-based sharpening of cryo-EM maps. Bioinforma Oxf Engl 36: 765–772. doi: 10.1093/bioinformatics/btz671

Rozov A, Khusainov I, El Omari K, Duman R, Mykhaylyk V, Yusupov M, Westhof E, Wagner A, Yusupova G. 2019. Importance of potassium ions for ribosome structure and function revealed by long-wavelength X-ray diffraction. Nat Commun 10: 2519. doi: 10.1038/s41467-019-10409-4

Scheres SHW, Chen S. 2012. Prevention of overfitting in cryo-EM structure determination. Nat Methods 9: 853–854. doi: 10.1038/nmeth.2115

Shao S, Murray J, Brown A, Taunton J, Ramakrishnan V, Hegde RS. 2016. Decoding Mammalian Ribosome-mRNA States by Translational GTPase Complexes. Cell 167: 1229–1240.e15. doi: 10.1016/j.cell.2016.10.046

Simonetti A, Guca E, Bochler A, Kuhn L, Hashem Y. 2020. Structural Insights into the Mammalian Late-Stage Initiation Complexes. Cell Rep 31: 107497. doi: 10.1016/j.celrep.2020.03.061

Sloan KE, Warda AS, Sharma S, Entian K-D, Lafontaine DLJ, Bohnsack MT. 2017. Tuning the ribosome: The influence of rRNA modification on eukaryotic ribosome biogenesis and function. RNA Biol 14: 1138–1152. doi: 10.1080/15476286.2016.1259781

Su H, Xu T, Ganapathy S, Shadfan M, Long M, Huang TH-M, Thompson I, Yuan Z-M. 2014. Elevated snoRNA biogenesis is essential in breast cancer. Oncogene 33: 1348–1358. doi: 10.1038/onc.2013.89

Tafforeau L, Zorbas C, Langhendries J-L, Mullineux S-T, Stamatopoulou V, Mullier R, Wacheul L, Lafontaine DLJ. 2013. The complexity of human ribosome biogenesis revealed by systematic nucleolar screening of Pre-rRNA processing factors. Mol Cell 51: 539–551. doi: 10.1016/j.molcel.2013.08.011

Taoka M, Nobe Y, Yamaki Y, Sato K, Ishikawa H, Izumikawa K, Yamauchi Y, Hirota K, Nakayama H, Takahashi N, et al. 2018. Landscape of the complete RNA chemical modifications in the human 80S ribosome. Nucleic Acids Res 46: 9289–9298. doi: 10.1093/nar/gky811

Tegunov D, Cramer P. 2019. Real-time cryo-electron microscopy data preprocessing with Warp. Nat Methods 16: 1146–1152. doi: 10.1038/s41592-019-0580-y

Valleron W, Laprevotte E, Gautier E-F, Quelen C, Demur C, Delabesse E, Agirre X, Prósper F, Kiss T, Brousset P. 2012a. Specific small nucleolar RNA expression profiles in acute leukemia. Leukemia 26: 2052–2060. doi: 10.1038/leu.2012.111

Valleron W, Ysebaert L, Berquet L, Fataccioli V, Quelen C, Martin A, Parrens M, Lamant L, de Leval L, Gisselbrecht C, et al. 2012b. Small nucleolar RNA expression profiling identifies potential prognostic markers in peripheral T-cell lymphoma. Blood 120: 3997–4005. doi: 10.1182/blood-2012-06-438135

van de Waterbeemd M, Tamara S, Fort KL, Damoc E, Franc V, Bieri P, Itten M, Makarov A, Ban N, Heck AJR. 2018. Dissecting ribosomal particles throughout the kingdoms of life using advanced hybrid mass spectrometry methods. Nat Commun 9: 2493. doi: 10.1038/s41467-018-04853-x

Watson ZL, Ward FR, Méheust R, Ad O, Schepartz A, Banfield JF, Cate JH. 2020. Structure of the bacterial ribosome at 2 Å resolution. eLife 9: e60482. doi: 10.7554/eLife.60482

Williams CJ, Headd JJ, Moriarty NW, Prisant MG, Videau LL, Deis LN, Verma V, Keedy DA, Hintze BJ, Chen VB, et al. 2018. MolProbity: More and better reference data for improved all-atom structure validation. Protein Sci Publ Protein Soc 27: 293–315. doi: 10.1002/pro.3330

Yu Y, Ji H, Doudna JA, Leary JA. 2005. Mass spectrometric analysis of the human 40S ribosomal subunit: native and HCV IRES-bound complexes. Protein Sci Publ Protein Soc 14: 1438–1446. doi: 10.1110/ps.041293005

Zhang D, Zhou J, Gao J, Wu R-Y, Huang Y-L, Jin Q-W, Chen J-S, Tang W-Z, Yan L-H. 2019. Targeting snoRNAs as an emerging method of therapeutic development for cancer. Am J Cancer Res 9: 1504–1516. eCollection2019

Zheng SQ, Palovcak E, Armache J-P, Verba KA, Cheng Y, Agard DA. 2017. MotionCor2: anisotropic correction of beam-induced motion for improved cryo-electron microscopy. Nat Methods 14: 331–332. doi: 10.1038/nmeth.4193

Zhou F, Liu Y, Rohde C, Pauli C, Gerloff D, Köhn M, Misiak D, Bäumer N, Cui C, Göllner S, et al. 2017. AML1-ETO requires enhanced C/D box snoRNA/RNP formation to induce self-renewal and leukaemia. Nat Cell Biol 19: 844–855. doi: 10.1038/ncb3563

Zivanov J, Nakane T, Forsberg BO, Kimanius D, Hagen WJ, Lindahl E, Scheres SH. 2018. New tools for automated high-resolution cryo-EM structure determination in RELION-3. eLife 7: e42166. doi: 10.7554/eLife.42166

